# A Novel TRPC6 Mutation Causes Autosomal Dominant FSGS

**DOI:** 10.1101/2025.02.11.637765

**Authors:** Rohan Bhattacharya, Xingrui Mou, Sophia M. Leeman, Alekshyander Mishra, Titilola D. Kalejaiye, Megan Stangl, Daniel Silas, Karen Soldano, C. John Sperati, Opeyemi Olabisi, Gentzon Hall, Samira Musah

## Abstract

FSGS is the most common primary glomerular lesion that causes kidney failure in the US. FSGS results from injury or loss of glomerular visceral epithelial cells (i.e. podocytes). Pathogenic variants in *TRPC6* can cause FSGS through dysregulated calcium conductance and associated disturbances in podocyte physiology. Here, we describe a 5-generation kindred with FSGS caused by a novel compound C-terminal TRPC6 mutation. Analysis of patient-specific iPSC-derived podocytes and glomerular capillary wall-on-a-chip systems revealed that the compound variants disrupt TRPC6 protein structure, intermolecular interactions, and membrane localization. Combined therapy with Sildenafil and Losartan ameliorated these disturbances compared to conventional immunosuppressive treatment.

## INTRODUCTION

Chronic kidney disease (CKD) is a progressive disease that can result in end-stage kidney disease (ESKD)^1^. Proteinuric kidney diseases are a leading cause of CKD and ESKD worldwide,^2^ and can be caused by podocyte injury or loss. Podocytes are an essential cellular component of the tripartite glomerular filtration barrier that form and maintain the filtration slit diaphragm (SD). Podocytes are highly specialized cells with limited replicative capacity; consequently, there has been considerable interest in developing rational, podocyte-sparing therapies for treating proteinuric kidney diseases^3^. Studies of familial forms of proteinuric kidney disease have identified pathogenic variants in more than 60 podocyte-expressed genes as the cause of focal segmental glomerulosclerosis (FSGS), the most common primary glomerular lesion that causes ESKD in the US^3^. Pathogenic variants in Transient Receptor Potential Cation Channel 6 (*TRPC6*) were first described as a cause of autosomal dominant FSGS in 2005^4^. To date, nearly 30 disease-causing *TRPC6* mutations have been documented, and dysregulated TRPC6 activity contributes to genetic and non-genetic forms of podocytopathy^5–7^. Pathogenic *TRPC6* mutations cause autosomal dominant disease with high penetrance^8–10^. Most TRPC6 podocytopathies are caused by gain-of-function (GOF) variants; however, one loss-of-function variant (G757D) has been associated with FSGS^11^. Dysregulation of TRPC6-mediated calcium conductance was shown to disrupt the podocyte actin cytoskeleton, overwhelm mitochondrial calcium buffering capacity, activate proapoptotic signaling, and induce podocyte dedifferentiation in various experimental models of glomerular disease^6^. Calcineurin inhibitors (CnIs) (i.e. cyclosporin A, voclosporin, and tacrolimus) have been the major class of immunosuppressive therapy for the treatment of TRPC6-associated nephropathy^12^; however, their long-term use can cause renal functional impairment and increase the risk of infectious complications^13,14^. Hence, an aggressive search for therapeutic alternatives has been long-standing. Advances in stem cell biology and bioengineering technology present new opportunities to precisely define the disease-relevant molecular mechanisms of TRPC6-induced podocyte injury in affected individuals and identify rational, podocyte-sparing therapeutics^15^.

Here, we report a 5-generation Northern European kindred (DUK40130) with biopsy-proven FSGS caused by a novel, compound, C-terminal *TRPC6* variant (p.L899H and p.P924T). An affected sibling pair from the family with non-nephrotic range proteinuria underwent clinical genetic evaluation following the identification of FSGS histology in the proband kidney biopsy. By electron microscopy, a dilated and highly fragmented endoplasmic reticulum (ER) system with numerous electron-dense ER inclusions was observed in podocytes. Similar findings were evident with high-resolution confocal microscopy, which revealed an accumulation of large podocin and TRPC6-enriched ER inclusions associated with disorganized TRPC6 and nephrin expression in iPSC-derived podocytes from both affected siblings. Assessment of podocyte barrier function using a bioengineered glomerular capillary wall-on-a-chip demonstrated significant albumin leakage with patient-derived podocytes relative to healthy controls. Treatment of iPSC-derived podocytes with an angiotensin receptor blocker (Losartan) and a calcineurin inhibitor (Tacrolimus) partially ameliorated albuminuria in the glomerular capillary wall-on-a-chip, recapitulating the effects of combination therapy in the affected siblings, but failing to reduce the formation of protein aggresomes in the ER. Combined therapy with the Phosphodiesterase 5 (PDE5) inhibitor Sildenafil and Losartan significantly improved podocyte barrier function and reduced albumin leakage in patient-derived podocytes. This combination therapy also significantly reduced podocin-rich aggresomes and restored TRPC6 and nephrin localization to podocyte arborizations.

## METHODS

### GENERATION OF PATIENT-SPECIFIC INDUCED PLURIPOTENT (iPS) CELLS

Protocols for the characterization of the patients and the creation of iPS cells were developed in accordance with the requirements of the institutional review board at Duke University Hospital. The kindred’s parents provided written informed consent. Details can be found in the Supplementary Appendix.

### PODOCYTE AND ENDOTHELIAL CELL DIFFERENTIATION

Details regarding podocyte and endothelial cell differentiation are provided in the Supplementary Appendix. Briefly, iPS cells were differentiated into mesoderm, then intermediate mesoderm cells, and finally into podocytes by activating TGFß, Wnt, Retinoic acid, and BMP7 signaling pathways ^16^. For each patient, the same iPS cells were also differentiated into lateral mesoderm and then endothelial cells by activating BMP4 and Wnt signaling^17^.

### COMPUTATIONAL MODELING OF TRPC6 PROTEIN STRUCTURE

ColabFold was utilized to make AlphaFold predictions of the structure of a single TRPC6 chain containing variants L899H and P924T^18^. The resulting structure was then comparatively analyzed against the reference PDB contained in the AlphaFold database utilizing the molecular graphics software UCSF ChimeraX^19^. The protein structure was analyzed for both localized and global changes. UCSF ChimeraX was also utilized to predict the structure of the TRPC6 four-chain multimer by superimposing Alpha Fold’s single-chain predictions onto the reference multimer contained in the UniProt database based on the matchmaker function. The multimer was then analyzed for changes in pore diameter utilizing the tape measure tool in UCSF ChimeraX. These predictions and analyses were repeated with the wildtype sequence as a negative control and six known mutations (P112Q, L395A, G757D, L780P, P924R, P924S) as positive controls (data not shown).

### GLOMERULAR CAPILLARY WALL-ON-A-CHIP DEVICE ENGINEERING

The organ-on-a-chip device was built with two channels separated by a biomimetic silk-based membrane^20^. Human iPS-derived intermediate mesoderm cells were seeded in the top channel and differentiated into podocytes in the presence of iPS-derived endothelial cells seeded in the bottom channel. The vascularized chips were maintained for at least nine days in culture under constant fluid flow and then analyzed using confocal and electron microscopy.

### STATISTICAL ANALYSES

We performed a Two-way Analysis of Variance (ANOVA) with Dunnett’s multiple comparison post hoc analyses with a family-wise alpha threshold of 0.05 and with individual mean difference computed for each comparison for Fig. 2E. We performed one-way ANOVA with Sidak’s multiple comparison post hoc analyses with a family-wise alpha threshold of 0.05 for Fig. 2G and Tukey’s multiple comparison post hoc analyses for Fig. 3F. We performed two-way ANOVA with Tukey’s multiple comparison post hoc analyses test for Fig. 3B-E and unpaired t-test for Fig. 3G.

## RESULTS

### MOLECULAR GENETIC ANALYSES

The DUK40130 proband (Duke 5, Female) and her affected twin sibling (Duke 7, Female) (**Fig. 1A**) were diagnosed with proteinuria at the age of 22 with dipstick proteinuria. Quantitation of urine protein-to-creatinine ratio (UPCR) at age 24 revealed 1.1 and 2.1 mg/mg, respectively. Both individuals started therapy with Losartan at age 24 and tacrolimus was added at age 25. The UPCR has remained at ∼ 0.8 - 1 mg/mg with normal eGFR (**Fig. 1B**). Exome analyses of Duke 5 and Duke 7 revealed two single-nucleotide substitutions (c.2696 T > A and c.2770 C > A) in *TRPC6*, which result in the pathogenic L899H and P924T variants of the C-terminal helical region of the protein (**Fig. 1C**). The variants are predicted to cause distortion of the C-terminal alpha helix, which was not observed in the reference or predicted WT structures (**Fig. 1D)**. The variants have not been documented together in the ∼200,000 individuals present in gnomAD and ClinVar databases. Computational modeling of the novel compound variants revealed greater tertiary structural changes (RMSD ∼19.2 Angstroms) than other known variants (RMSD ∼17.2-18.9 Angstroms), without a significant alteration of the channel pore diameter (WT TRPC6 diameter = 9.71 Angstroms, predicted WT TRPC6 diameter = 5.67 Angstroms, and TRPC6^899H,924T^ diameter = 5.85 Angstroms) (**Fig. 1E, Fig S1A**). This distortion of the C-terminal alpha helix can possibly be explained by the removal of the helix-breaking proline at position 924^21,22^ (**Fig 1D**). Biopsy of Duke 5 revealed focal glomerular sclerosis and hyalinosis by light microscopy (**Fig. 1F)**. Electron microscopy demonstrated podocyte foot process (FP) effacement and marked disruption of podocyte ER architecture associated with numerous electron-dense ER inclusions (**Fig. 1G**. TRPC6 is a major component of the podocyte SD complex, where it associates with other essential SD proteins, including nephrin and podocin^23^. Podocin and TRPC6 interact at their respective C-termini. This region of protein-protein contact operates as a molecular switch to negatively modulate TRPC6 activity^24^. Given the functional importance of this known intermolecular interaction, the presence of the variants within the region, and the unique presence of podocyte ER inclusions associated with the variants, we hypothesized that the L899H/P924T variants may physically disrupt Nephrin-Podocin-TRPC6 complex formation, leading to trafficking disturbances and ER congestion.

**Figure 1.**
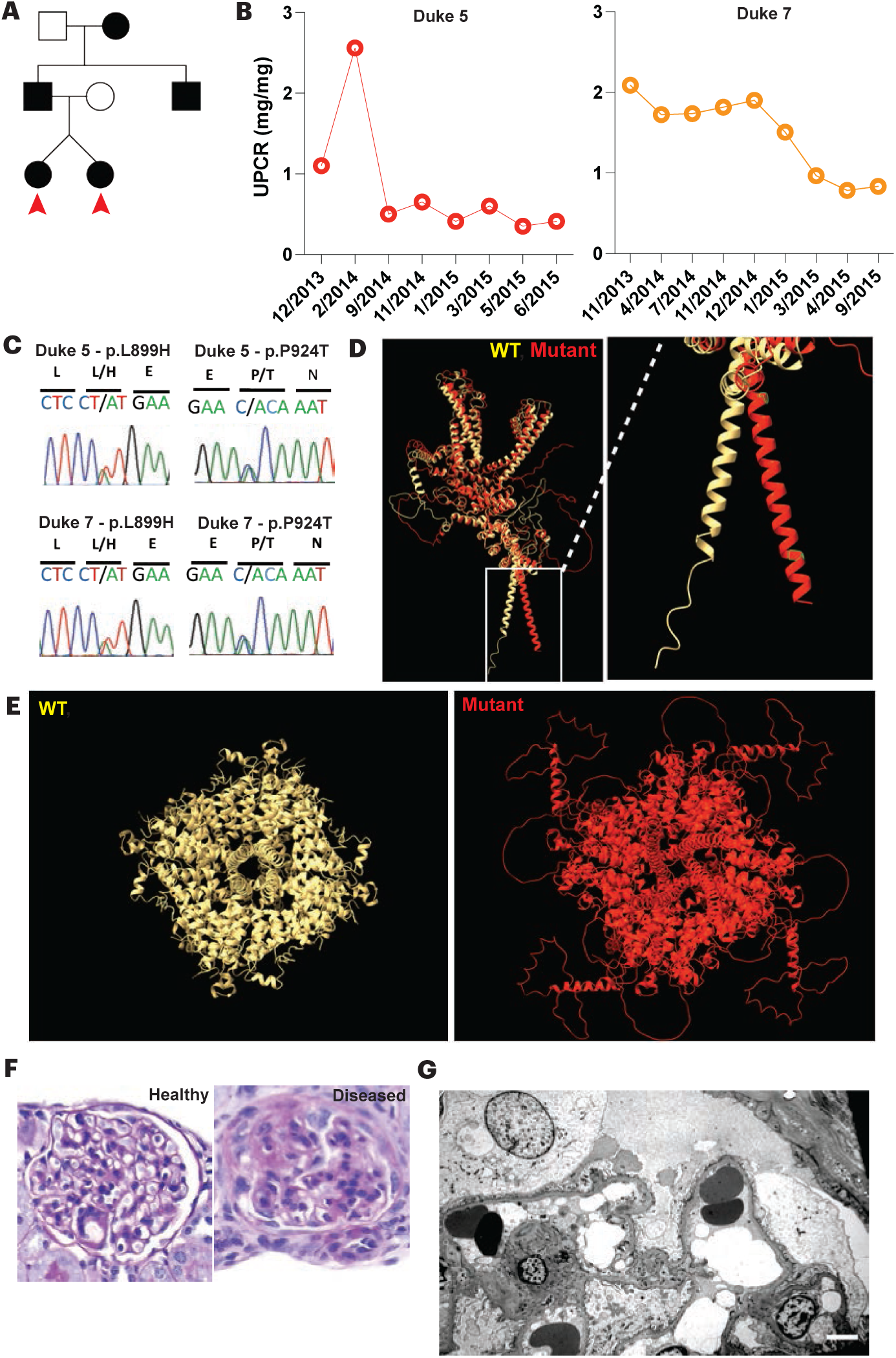
Twins with evidence of novel compound TRPC6 mutation-associated glomerulopathy. Panel A shows the family pedigree of the twin Northern European kindred (DUK40130) with biopsy-proven Focal Segmental Glomerulosclerosis (FSGS). Panel B shows the urinary protein creatine ratio (UPCR) in Duke 5 and Duke 7. Panel C shows the sequence of the TRPC6 C terminus. Both patients had a novel compound, C-terminal TRPC6 mutation (p.L899H and p.P924T). Panel D shows the computationally generated superimposed images of the WT reference and computationally predicted mutant single-chain TRPC6 proteins with a focus on the C terminal region of the protein. Panel E shows the pore architecture of the WT and the mutant TRPC6 protein (WT TRPC6 diameter = 9.71 Angstroms, predicted WT TRPC6 diameter = 5.67 Angstroms, and TRPC6^899H,924T^ diameter = 5.85 Angstroms). Panel F shows a hematoxylin and eosin staining of Duke 5’s kidney biopsy with focal glomerular sclerosis and hyalinosis. Panel G shows an electron micrograph of Duke 5’s biopsy with dilated ER with electron-dense ER congestion.

### SLIT DIAPHRAGM PROTEIN LOCALIZATION IS ALTERED IN PATIENTS’ PODOCYTES

Podocyte differentiation from the Duke 5 and Duke 7 iPS cells revealed large podocin-enriched ER inclusions associated with significantly reduced TRPC6 and nephrin expression (**Fig. 2A, Fig. S1B and C**). There is significant disruption of nephrin and podocin colocalization in Duke 5 and Duke 7 podocytes relative to healthy controls as confirmed by Mander’s correlation coefficient (**Fig. S1D**). Nephrin and podocin are essential components of the filtration SD that also facilitate the assembly of pro-survival signaling assemblies and cytoskeletal regulatory networks at the glomerular filtration barrier (GFB)^7^. Given the established role of TRPC6 in regulating podocyte actin-cytoskeleton^9,25^, we questioned how the compound L899H and P924T variants might impact nephrin-podocin-TRPC6 subcellular localization to terminal podocyte arborizations. Immunostaining of healthy control podocytes revealed colocalized punctate expression of nephrin, podocin, and TRPC6 in terminal podocyte arborizations. Conversely, the colocalized expression of these markers was significantly reduced in Duke 5 and Duke 7 podocytes (**Fig. 2B, S1E**).

**Figure 2.**
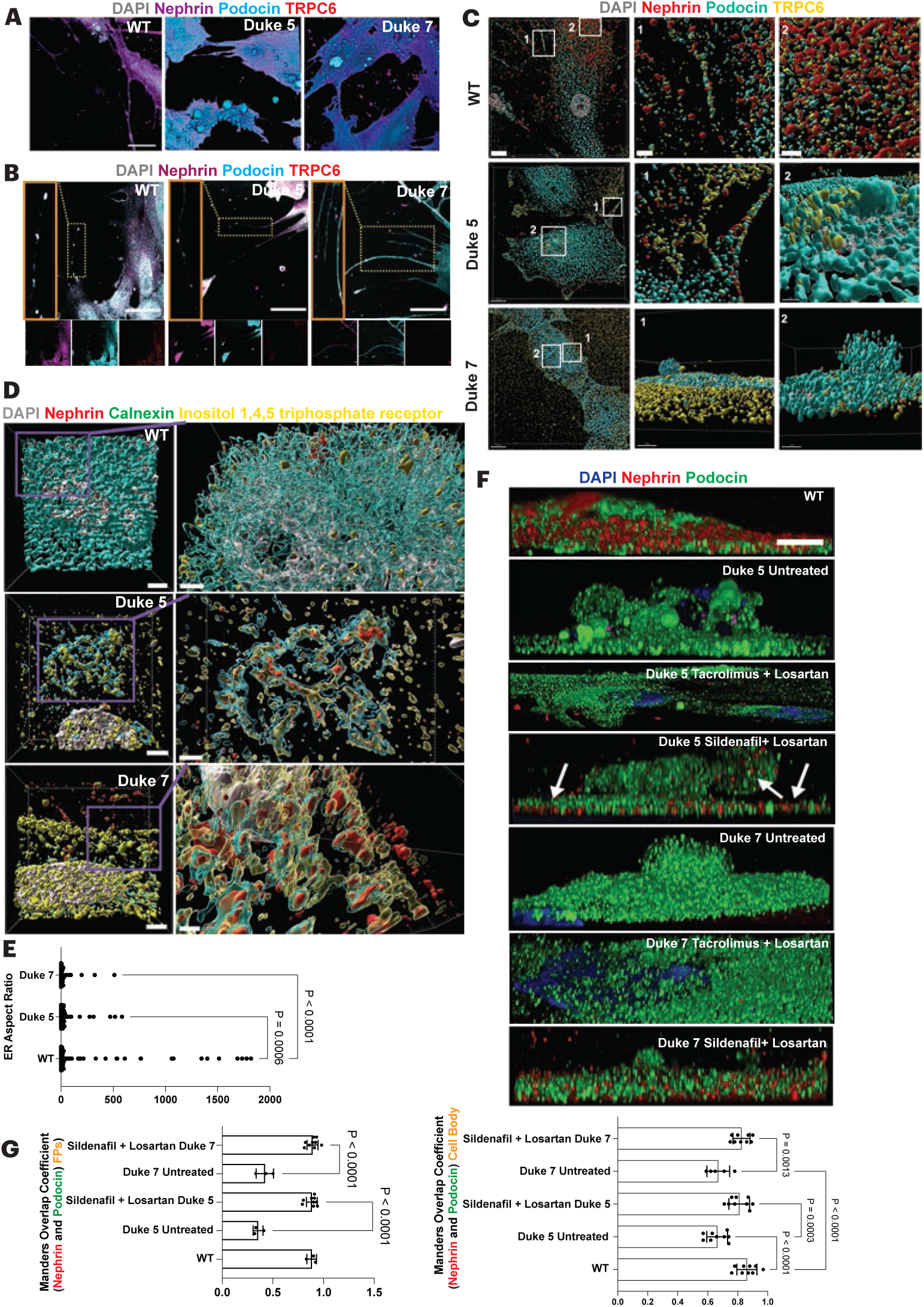
Podocyte-derived from patient-specific human iPS cells demonstrate altered expression of podocyte-lineage markers and dilated endoplasmic reticulum. Panel A shows the immunofluorescence of patient-specific human induced pluripotent stem (iPS) cell-derived podocytes’ cell bodies labeled with nephrin, podocin, and TRPC6. Panel B shows the immunofluorescence of patient-specific iPS cell-derived podocytes’ foot processes labeled for nephrin, podocin, and TRPC6 expression. Panel C shows the distribution of nephrin, podocin, and TRPC6 in the podocytes’ cell bodies and foot processes. Panel D shows healthy patient podocytes immunostained for nephrin, calnexin, and ionositol 1,4,5 triphosphate receptors. Panel E shows the quantification of the ER aspect ratio based on calnexin expression in podocytes. Panel F shows the immunofluorescence of treated and untreated patient-specific podocytes immunostained for nephrin, podocin, and TRPC6. Panel G shows the Mander’s overlap coefficient for nephrin and podocin in the podocytes’ cell bodies and foot processes before and after treatment with sildenafil and losartan. Panels E and G represent three biologically independent experiments, and the data is defined as ± standard error of the mean. Scale bars: Panel A, 20 µm; Panel B, 20 µm; Panel C, column 1 images 15 µm, column 2 and 3 images 5 µm; Panel D column 1, 4 µm; column 2, 2 µm; Panel F, 5 µm.

Next, we investigated how SD proteins interact spatially within the cell body and terminal podocyte arborizations. A 3D rendering of high-resolution confocal images revealed a uniform distribution of the nephrin-podocin-TRPC6 complexes throughout the cytoplasm and in the cell body of WT podocytes, highlighting the importance of these protein-protein interactions in podocyte morphological phenotype and molecular physiology. Punctate localization of nephrin-podocin-TRPC6 in podocyte arborizations was also observed. By contrast, this spatial patterning was disrupted in Duke 5 and Duke 7 podocytes, consistent with an impairment in SD complex formation in the diagnostic biopsy (**Figs. 1G and 2C**). Immunoblots from patient-specific differentiated podocytes also revealed significant downregulation of nephrin, TRPC6, and synaptopodin (**Fig. S2A, B**) with massive podocin-enriched aggresomes in the ER (**Fig. S2C**).

To further explore potential disruption in trafficking or endosomal processing of the SD proteins, we evaluated whole cell lysates from Duke 5 and Duke 7 podocytes by immunoblot. We demonstrated the presence of high molecular weight podocin aggresomes were almost nonexistent in the WT. Evaluation of the ER morphology by high-resolution confocal microscopy demonstrated a regular perinuclear distribution of ER tubules in healthy controls, while the ER tubules in mutant podocytes exhibited a fragmented and attenuated appearance with mislocalized nephrin expression and high inositol 1,4,5 triphosphate receptor indicating calcium efflux from the ER (**Fig. 2D, E**). We then examined the expression of late endosome and lysosomal biogenesis markers LAMP1 and RAB7, respectively. While LAMP1 vesicle number remained similar in mutant and healthy podocytes (**Fig. S2D**), RAB7 expression was significantly downregulated in the mutant podocytes, suggesting that Duke 5 and Duke 7 podocytes may exhibit reduced lysosomal biogenesis, leading to uncontrolled protein accumulation and ER congestion. (**Fig. S2E**).

### SILDENAFIL AND LOSARTAN COTREATMENT REDUCE PODOCIN-RICH AGGRESOMES AND RECRUIT NEPHRIN

Next, we sought to evaluate non-immunosuppressive candidate therapies that could improve podocyte SD marker localization and resolve podocin-enriched aggresomes. To recapitulate patient clinical management, we first treated podocytes with tacrolimus and losartan (TL). Combination therapy with these agents did not restore nephrin-podocin colocalization (**Fig. 2F**). We also tested the effects of the selective TRPC6 blocker, BI749327, which has been shown to suppress TRPC6-mediated cation conductance and inhibit activation of Nuclear Factor of Activated T-cell (NFAT) in HEK cells overexpressing disease-causing gain-of-function TRPC6 mutations (i.e. P112Q, M132T, R157Q, R895C, and R895L)^26^. BI749327 failed to reduce podocin aggresomes and improve nephrin-podocin-TRPC6 colocalization in Duke 5 and Duke 7 podocytes, suggesting that the formation of the ER aggresomes was unrelated to aberrations in TRPC6-mediated cation conductance (**Fig. S3A**). We also tested patient-derived podocytes with sparsentan, a dual endothelin-angiotensin II receptor antagonist that can suppress TRPC6-mediated calcium influx with preservation of podocyte nephrin and podocin expression^27^. Sparsentan also failed to reduce podocin-rich aggresomes or to reestablish nephrin-podocin colocalization (**Fig. S3B**), highlighting again that the formation of ER protein aggresomes in the mutant podocytes may be independent of dysregulated TRPC6-mediated cation conductance^28^. When Duke 5 and Duke 7 podocytes were treated with the Phosphodiesterase 5 (PDE5) inhibitor, Sildenafil^29,30^ in combination with Losartan (SL), we observed a significant reduction in podocin-rich aggresomes and complete restoration of nephrin-podocin colocalization (**Fig. 2F, G**) which has important implications in bridging podocyte-podocyte junctions. Although PDE5 inhibitors have been shown to downregulate TRPC6 activity^29^, it is likely that the beneficial effect of restoring podocyte proteostasis are pleiotropic and independent of the ability to suppress aberrant TRPC6 activity^31^.

### SILDENAFIL AND LOSARTAN AMELIORATE ALBUMINURIA IN AN ENGINEERED GLOMERULAR CAPILLARY WALL-ON-A-CHIP

To explore the effects of therapy on podocyte barrier functions, we engineered a vascularized glomerular capillary wall-on-a-chip platform to recapitulate the GFB using patient-derived isogenic iPS cells (**Fig. 3A**). The tripartite glomerular filtration barrier is comprised of endothelium on the luminal surface of the glomerular basement membrane and terminally differentiated podocytes on the opposing urinary surface. Podocytes, with their interdigitating foot processes, provide the primary support for the filtration slit diaphragm, a heteroporous, zipper-like assembly of cadherin and immunoglobulin-like proteins that possess molecular charge and molecular size selectivity properties for ultrafiltration^32^. After differentiation on the microfluidic chip, immunostaining of mutant podocytes revealed significantly elevated levels of podocin and TRPC6 and reduced levels of nephrin and CD2AP compared to WT (**Fig. S4**). Treatment with TL or SL, significantly reduced podocin and TRPC6 aggregation in Duke 5 and Duke 7 podocytes (**Figs. 3B-E, S5**). However, Duke 5 and Duke 7 podocytes treated with SL exhibited higher Nephrin and CD2AP levels as compared with their TL-treated counterparts (**Figs. 3C, 3E, S5**). These findings demonstrate that SL more effectively restores the pattern of SD protein localization than TL in mutant podocytes. Next, we examined albumin excretion by quantifying albumin filtration into the “urinary” channel effluent of the microfluidic chips. We demonstrated that the effluent from Duke 5 chips had significantly higher albumin levels relative to controls, consistent with the clinical manifestation of the disease in the affected siblings (**Fig. 3F**). The mutant podocytes also demonstrated reduced albumin sequestration from the filtrate (**Fig. S6A**). Treatment with both SL and TL reduced albumin leakage (**Fig. 3F**) in the Duke 5 chips; however, SL was superior to TL in reducing albumin excretion. Albumin leakage in Duke 7 chips remained similar to WT chips after SL treatment, while TL-treated Duke 7 chips demonstrated increased albumin excretion (**Fig. 3F**). Subsequent TEM imaging of untreated Duke 5 and Duke 7 podocytes on the microfluidic chips revealed fragmented ER tubules with ER inclusions. In SL-treated patients’ podocytes, ER aggresomes were significantly reduced (**Fig. 3G**), and ER morphology was markedly improved (**Fig. S6B**), consistent with a beneficial effect of improved proteostasis on podocyte barrier function.

**Figure 3.**
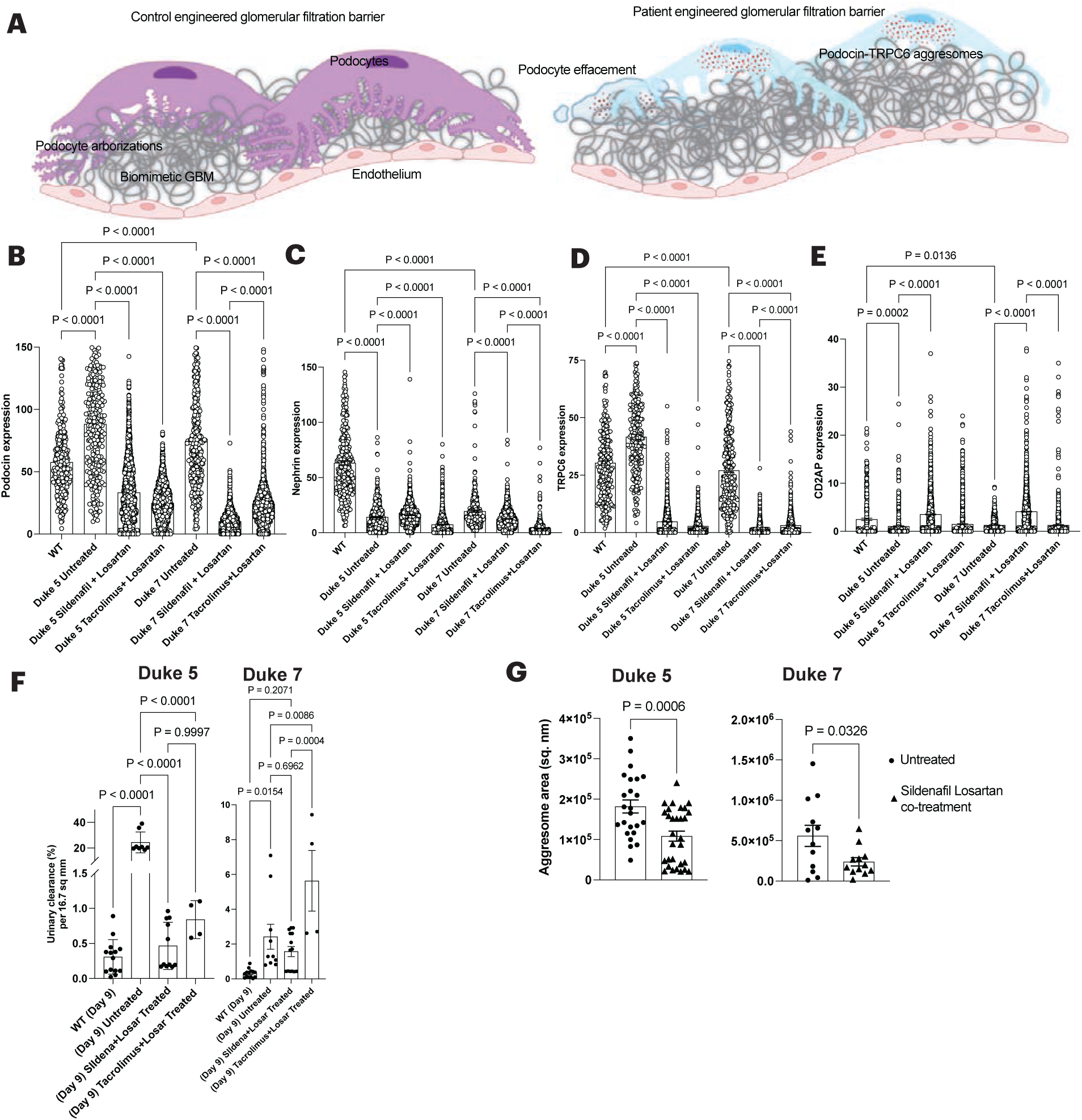
Sildenafil and Losartan co-treatment reduces aggresomes and increases Nephrin and CD2AP expression in the patient-specific glomerular-capillary wall on a chip. Panel A shows the schematic representation of an engineered glomerular capillary wall-on-a-chip with patient-specific iPS cell-derived podocytes and endothelial cells. The protein aggresomes are shown in the mutant organ chip. Podocyte lineage-specific marker expression in the glomerular capillary wall-on-a-chip was measured for Panel B, podocin; Panel C, nephrin; Panel D, TRPC6; Panel E, CD2AP. Panel F shows the urinary clearance % quantified from the urinary channel after perfusing the chips with fluorescently labeled albumin for 6 hours. Panel G shows protein aggresome size in podocytes before and after sildenafil and losartan combinatorial therapy, as represented in Figure S6B. Panels B to G represent three biologically independent experiments, and the data is defined as ± standard error of the mean. Each data point represents signal intensity emanating from the immunostained chips/pixel^2^ as represented in Figure S5 across an area of ∼75 µm.

## DISCUSSION

In this report, we describe novel compound C-terminal TRPC6 variants that induce podocyte injury via alteration of SD protein expression and targeting, disruption of SD complex assembly, and ER congestion/fragmentation. These novel features of TRPC6-associated podocyte injury and dysfunction were amenable to therapeutic correction using combined ARB and PDE5 inhibition, but were not ameliorated by treatment with tacrolimus or other agents targeting TRPC6-mediated calcium conductance. Although the precise mechanisms of the therapeutic benefit are yet to be determined, it is well-known that PDE5 inhibitors can ameliorate dysregulated TRPC6 activity in various cell types^29,30^. Inhibition of PDE5 catalytic activity prevents the hydrolytic conversion of cyclic guanosine monophosphate (cGMP) to the inactive metabolite, 5’ GMP^33^.cGMP is an essential activator of Protein Kinase G, a member of the AGC kinase family that has been shown to inhibit TRPC6 activity and expression via direct phosphorylation of the threonine residue 69 in podocytes^34^. PKG has also been shown to positively regulate the proteasome-mediate degradation of misfolded proteins, to promote proteostasis^35^, to enhance autophagic flux and to reduce protein aggregation^36^. PDE5 inhibitors have also been shown to upregulate the expression of peroxisome proliferator-activated receptor γ (PPARγ)^30^. PPARγ is a nuclear transcription factor and positive regulator of autophagy and lysosomal biogenesis via suppression of mTOR activity. Previous work showed that Sildenafil suppressed podocyte TRPC6 expression through the recruitment of PPARγ to the TRPC6 promoter, demonstrating a role for the PPARγ as a transcriptional repressor^30^. Although our studies do not delineate the contributions of PKG and PPARγ signaling to the amelioration of podocyte injury caused by the compound L899H/P924T variants, it is possible that the known cytoprotective actions of these molecules on proteostasis could underlie the beneficial effects we observe on podocyte morphology and function. Although no direct influence of ARBs on proteostasis have been defined, the potential for additive or synergistic benefit from combined therapy on podocyte function cannot be excluded by our studies and warrant further investigation.

Dysregulated TRPC6 activity can significantly contribute to many non-genetic forms of podocytopathy, such as diabetic nephropathy, membranous nephropathy, and minimal change disease^37^. Consequently, deciphering the molecular mechanisms of TRPC6-mediated podocytopathy is a subject of intense scientific and clinical/translational interest. While the precise mechanisms of TRPC6-mediated injury across the spectrum of glomerular disease are largely undefined, aberrant calcium transport has been identified as a key feature of the disease-relevant physiologic disturbances in podocytes. In this study, we reported novel compound C-terminal TPRC6 variants (L899H/P924T) that induce the accumulation of podocin-enriched ER aggresomes, disrupts ER morphology, and impairs nephrin-podocin-TRPC6 trafficking to actin-based, terminal podocyte arborations. This is the first study to identify these features of TRPC6-mediated podocyte injury and explore the therapeutic utility of targeting TRPC6-induced alterations in podocyte proteostasis and suggest various pathogenic mechanisms for the effects of TRPC6 mutations. Our findings expand the spectrum of TRPC6-mediated podocyte cellular injury and provide further evidence of the pleiotropic benefits of PDE5 inhibitors. What remains unknown are the specific pathomechanisms of L899H/P924T on TRPC6 interactions with nephrin and podocin, the role of impaired proteostasis in other forms of TRPC6-mediated podocytopathy, and the molecular underpinnings of PDE5 inhibitor-induced podocyte cytoprotection.

## SUPPLEMENTARY APPENDIX

### Patient-specific cell isolation

Whole blood was collected for Duke 5 and Duke 7 using EDTA blood collection tubes. Peripheral blood mononuclear cells (PBMCs) were isolated by density gradient centrifugation with SepMate-50 PBMC Isolation Tubes (StemCell Technologies, 85450) used in conjunction with the density gradient medium Lymphoprep (StemCell Technologies, 07811) per manufacturer’s protocol. PBMCs were then washed with DPBS with 2% FBS (StemCell Technologies, 07905) and cryopreserved in CryoStor CS10 (Stemcell Technologies, 07930) for long-term storage in liquid nitrogen.

### Human iPS cell generation

To generate iPSCs derived from Duke 5 and Duke 7, PBMCs were thawed gently at 37°C and diluted 1:10 into DPBS with 2% FBS to remove cryopreservation solution. PBMCs were resuspended in StemSpan SFEM II (StemCell Technologies, 09655), StemSpan Erythroid Expansion Supplement (1X) (Stemcell Technologies, 02692), Primocin (0.2%) (InvivoGen, ant-pm-1), and Penicillin-Streptomycin (1%) (Gibco, 15140122) and plated on a 6-well plate. PBMC media was changed every other day. After seven days PBMCs were collected and counted. 30,000 PBMCs were transduced (Day 0) using CytoTune-iPS 2.0 Sendai Reprogramming Kit (ThermoFisher Scientific, A16517) prepared in PBMC Media (without Primocin) following manufacturer’s protocol. Transduced cells were seeded into one well each of a 48-well plate and incubated for 24 hours in a 37°C incubator at 5% CO2. The media was changed into PBMC Media to remove Sendai virus and cells were incubated for an additional 48 hours. A feeder layer of mouse embryonic fibroblasts (MEFs) (Gibco, A24903) was prepared on Geltrex LDEV-Free, hESC-Qualified, Reduced Growth Factor Basement Membrane Matrix (Gibco, A1569601)-coated 12-well plates in MEF Media: DMEM (Gibco, 11965092), human Embryonic Stem Cell-qualified FBS (10%) (Gibco, 16141061), MEM Non-Essential Amino Acids Solution (1mM) (Gibco, 11140050), Beta-mercaptoethanol (55uM), Penicillin-Streptomycin (1%) (Gibco, 15140122). On Day 3, the transduced PBMCs were collected, resuspended in iPSC Media: DMEM F12 + GLutamax (Gibco, 10565018), Knockout Serum Replacement (20%) (Gibco, 10828010), MEM Non-Essential Amino Acids Solution (1X) (Gibco, 11140050), Penicillin-Streptomycin (1%) (Gibco, 15140122), Beta-mercaptoethanol (55uM) (Gibco, 21985023), Human FGF-basic (10ng/mL) (Peprotech, 100-18B), and seeded onto the MEF-coated plates. The freshly seeded plate was centrifuged at 210-300 RCF for five minutes in order to encourage cell-to-cell adhesion between the PBMCs and MEFs. Media was changed with fresh iPSC Media every other day or as needed. Between Days 19-21 robust iPSC colonies were dissociated from the MEFs with Versene Solution (Gibco, 15040066), incubated for two minutes at 37°C, picked via suction with a P200 pipette tip, and resuspended in StemFlex Basal Medium plus StemFlex Supplement (1X) (Gibco, A3349401), Penicillin-Streptomycin (1%) (Gibco, 15140122), and rho-kinase inhibitor Y-27632 dihydrochloride (0.1%) (Selleck Chem, S1049), and seeded into individual wells of a 12-well, Vitronectin (VTN-N) Recombinant Human Protein, Truncated (ThermoFisher Scientific, A14700) coated plate. Up to five clones per sample were selected. After 24 hours cells had adhered to the plate and media was changed to remove Y-27632. Cells were passaged such that confluence was maintained between 30 - 70% and media was changed every other day or as needed. Y-27632 was used in StemFlex media to passage cells and removed 24 hours after each passage. iPS cells morphology was monitored and assessed daily to ensure cells maintained compact structures with distinct borders. Ultimately, two clones per sample were selected for further maturation. After no fewer than 10-15 passages, iPS cells were considered to be stable and ready for quality assessments. Two iPSC clones from each sample (Duke 5 and Duke 7) were column-purified with anti-TRA-1-60 microbeads (Miltenyi Biotec, Inc., 130-100-832), passaged and plated on a 6-well VTN-N coated plate for expansion, along with two wells of a 96-well coated plate for assessment of iPS cells via immunocytochemistry. The Pluripotent Stem Cell 4-Marker Immunocytochemistry Kit (Invitrogen, A24881) was used to detect SOX2, TRA-1-60, OCT4, and SSEA4, known expression factors of mature iPSC cells. A cell pellet was collected and KaryoStat assay (Thermo Fisher Scientific, 905403) was performed to ensure no chromosomal abnormalities were introduced through the iPSC derivation process. Human iPS cells that passed quality control assessments were frozen in CryoStor CS10 and stored in liquid nitrogen until ready for downstream use.

### iPS cell culture

Patient-specific iPS cells (WT, Duke 5, and Duke 7) were propagated on 6-well plates coated with Matrigel Matrigel (VWR; 75796-278) by using mTeSR1 (StemCell Technologies; 85870) medium without antibiotics. The cells were split every 5 to 6 days (or until 75% confluency is reached) by treatment with Accutase (Thermo/Life Technologies; A1110501). The cell lines were propagated in a 37 °C incubator with 5% CO2. The cell lines were routinely tested for mycoplasma and the cells were found to be free of mycoplasma contamination (Mycoplasma PCR Detection Kit from abm. G238).

### Differentiation of mesoderm, IM, and podocytes

For directed differentiation of Duke 5 and Duke 7 iPS cells towards mesoderm, iPS cells were carefully dislodged with enzyme-free dissociation buffer (Gibco, 13150-016) and centrifuged once at 200 g for 5 mins in advanced DMEM/F12 (Gibco; 12634010) to remove residual Matrigel or stem cell culture media components. The supernatant was aspirated, and the cells were resuspended in mesoderm differentiation media. Cells were seeded in a laminin 511-E8 (Takara T304)-coated 12 well plate. The mesoderm differentiation media comprised of 100 ng/ml activin A (Thermo Fisher Scientific, PHC9561), 3 μM CHIR99021 (Stemgent, 04-0004), 10 μM Y27632 (TOCRIS, 1254), and 1× B27 serum-free supplement (GIBCO, 17504044) dissolved in DMEM/F12 with GlutaMax (GIBCO, 10565018)^16^. After 2 days of differentiation, the media was replaced with Intermediate Mesoderm Differentiation Media. The Intermediate Mesoderm Differentiation Media comprised of 100 ng/ml BMP7 (Thermo Fisher Scientific, PHC9541), 3 μM CHIR99021 (Stemgent, 04-0004), and 1× B27 serum-free supplement (GIBCO, 17504044) dissolved in DMEM/F12 with GlutaMax (GIBCO, 10565018)^16^. After 14 days of differentiation, Intermediate Mesoderm cells were trypsinized with 0.25% trypsin-EDTA (Gibco; 25200-056) and cells were replated on freshly prepared Laminin 511-E8 coated 12 well plates at a density of 100,000 cells/well with Podocyte Differentiation Medium. The Podocyte Differentiation Medium comprised of 100 ng/ml BMP7, 100 ng/ml activin A, 50 ng/ml VEGF (Thermo Fisher Scientific, PHC9391), 3 μM CHIR99021, 1× B27 serum-free supplement, and 0.1 μM all-trans retinoic acid (Stem Cell Technologies, 72262) dissolved in DMEM/F12 with GlutaMax (GIBCO, 10565018)^16^. The cells were cultured for 5 days with regular media change.

### Drug treatment

iPS-derived podocytes were maintained in CultureBoost-R for one day after the completion of the podocyte differentiation. For drug treatment, drugs (dissolved in DMSO) were added to CultureBoostR, mixed well, and introduced to the cells. DMSO served as a control. The drugs and their concentrations used in this study were as follows: Tacrolimus (MedchemExpress, HY-13756A, 1 µM), BI749327 (MedchemExpress, HY-111925, 500 nM), Sparsentan (MedchemExpress, HY-17621, 10 µM), Sildenafil (MedchemExpress, HY-15025,1 µM), and Losartan (MedChemExpress #HY-17512/CS-2116, 10 µM).

### Differentiation of endothelial cells

WT, Duke 5, and Duke 7 iPS cells were harvested with Accutase (Thermo/Life Technologies; A1110501) and reseeded on Matrigel-coated 6-well plates at a cell density of 420,000 cells/well. After 1 day of incubation, Lateral Mesoderm Differentiation Media was added to the wells. The Lateral Mesoderm Differentiation Media comprised of N2B27 media, which contains neurobasal media (Invitrogen, 21103049) and DMEM/F12 glutamax (Invitrogen) in 1:1 ratio with N2 (100×) (GIBCO, 21103049) and B27 without Vitamin A (GIBCO, 12587010), 8 µM CHIR99021 and 25 ng/mL hBMP4 (VWR International LLC, 10273-372) for 3 days. On Day 4, media was replaced with Endothelial cell Differentiation Media for an additional 3 days. The Endothelial cell Differentiation Media comprised of StemPro-34 SFM media (GIBCO, 10639011) supplemented with Glutamax (GIBCO, 35050061) in 100:1 ratio, 2 µM forskolin (Abcam Inc., ab120058), and 200 ng/mL VEGF165. Cells were fed daily, and conditioned media was collected for endothelial expansion media preparation. On Day 7, differentiated cells were dislodged with cold Accutase treatment for 5 mins and MACS-sorted (MidiMACS™ Separator) to harvest CD144^+^ and CD31^+^ cell populations. Purified cell populations were expanded in Endothelium Maintenance Media comprised of conditioned media diluted at 1:1 ratio with StemPro-34 SFM supplemented with 2 µg/mL heparin (STEMCELL Technologies, 07980). Media was replenished every other day or until the conditioned media was depleted. For continued expansion beyond the first passage, the cells were fed with viEC maintenance media, which comprised of StemPro-34 supplemented with 10% HI-FBS (Invitrogen, 10082147), 2 µg/mL heparin, and 50 ng/mL VEGF165^17^.

### Western Blot Analysis

WT, Duke 5, and Duke 7 podocytes were washed with cold PBS (Gibco; 14190144) on Day 6 and lysed on ice in RIPA buffer (Millipore/Sigma; R0278-500ML) supplemented with PhosSTOP phosphatase inhibitors (Millipore/Sigma; 04906837001) and complete EDTA-free protease inhibitor cocktail (Millipore/Sigma; 4693132001). One tablet of each PhosSTOP phosphatase inhibitor and complete EDTA-free protease inhibitor was added in 10 mL RIPA Buffer. Protein samples were separated by SDS-PAGE using Mini-PROTEAN TGX Stain-Free Precast 4-15% Gels (Bio-rad; 4568083) and transferred onto a PVDF membrane (Bio-rad; 1704157) using a Trans-blot Turbo semi-dry transfer system (Biorad; 1704150). Membranes were blocked with 5% Blotto (ChemCruz; sc-2324) in Tris-buffered saline with Tween 20 (TBST), and immunoblotting was carried out according to standard procedure. Primary antibodies were always diluted in TBST supplemented with 5% Blotto and incubated for 1 h at room temperature on a platform rocker. Primary antibodies used for Western blot were guinea pig Anti-Nephrin antibody (ARP; #GPN02, 1:500), rabbit Anti-Podocin (Abcam, mAb #ab50339, 1:1000), mouse Anti-Synaptopodin (D-9) (SantaCruz, mAb #sc-515842, 1:1000), mouse Anti-CD2AP (B-4) (SantaCruz, mAb #sc-25273, 1:1000), mouse Anti-WT1 (6F-H2) (Millipore Sigma #05-753, 1:250), mouse Anti-GLEPP1 (B-6) (SantaCruz, mAb #sc-365354, 1:1000), and rabbit Anti-GAPDH (Millipore Sigma; #ABS16, 1:10000). Secondary antibodies used in this study included HRP-conjugated goat anti-mouse (Cell Signaling; #7076, 1:5000), HRP-conjugated goat anti-rabbit (Cell Signaling; #7074, 1:10000), HRP-conjugated goat anti-guinea pig polyclonal (ARP; #90001, 1:1000), and HRP-conjugated donkey anti-goat polyclonal (R&D Systems, #HAF109, 1:1000). Secondary antibodies were also diluted in TBST supplemented with 5% Blotto and incubated for 1 h at room temperature on a platform angle rocker. Chemiluminescence was detected using the Super Signal West Femto kit (Thermo; 34094). Signal intensities were analyzed using a ChemiDoc imager (Bio-rad).

### Immunostaining and Microscopy

WT, Duke 5, and Duke 7 podocytes were washed with cold PBS (Gibco; 14190144) on Day 6 and were fixed with 4% paraformaldehyde (Sigma-Aldrich, #16005) in dPBS followed by permeabilization with 0.125% triton X-100 in dPBS for 5 min. Cells were blocked by incubation with a solution of 1% BSA and 0.125% triton X-100 in dPBS (blocking buffer) for 30 min on ice. The cells were incubated with primary antibodies in a permeabilization buffer overnight at 4 °C. The cells were washed with permeabilization buffer 3-4 times and incubated with secondary antibodies conjugated to either Alexafluor-488 (Life Technologies; #A21202 1:1000), Alexafluor-594 (Life Technologies; #A21203 1:1000) or Alexafluor-700 (Life Technologies; #A21038 1:1000) in permeabilization buffer for 1 h at room temperature. Following incubation, the cells were washed three times with permeabilization buffer and counterstained with 4′,6-diamidino-2-phenylindole (DAPI) (Invitrogen #D1306). The primary antibodies used included guinea pig Anti-Nephrin (ARP; #GPN02, 1:500), rabbit Anti-Podocin (Abcam, mAb #ab50339, 1:1000), rabbit Anti-TRPC6 (Abcam, pAb #ab228771) and mouse Anti-CD2AP (B-4) (Santa Cruz, #sc-25272), Calnexin (C5C9) Anti-Rabbit (Cell Signaling, mAb #2679), LAMP1 (D2D11) XP Anti-Rabbit (Cell Signaling, mAb #9091), RAB7 (D95F2) XP and Anti-Rabbit mAb (Cell Signaling, mAb #9367). Immunofluorescence images were captured using the Zeiss 780 Upright confocal microscope equipped with 63x/1.4 Oil Zeiss Plan-Apochromat 44 07 62 (02) WD 0.19 mm objective. Data was analyzed using the Zen 2.3 Black software.

### Generation of glomerular-capillary-on-a-chip

Biomimetic silk fibroin chips^20^ were treated with oxygen plasma at 50 W and 0.8 mbar for 60 s using a plasma asher (Emitech, K-1050X) to activate the membrane for surface adsorption of laminin-511 solution (25 μg/ml; Biolamina, LN511), followed by incubation at 37°C overnight. The next day, the channels were washed with advanced DMEM/F12 (Gibco; 12634010), and endothelial cells were seeded on the bottom channel. After 4 h incubation, isogenic Intermediate mesoderm cells were seeded on the top channel and incubated overnight at 37 °C. The next day, the top channel was flushed with Podocyte Differentiation Medium, and the bottom was flushed with viEC medium. The cells were then continuously perfused with their respective cell-culture media using an Ismatec IPC-N digital peristaltic pump (Cole-Parmer) at a volumetric flow rate of 246 μl h^−1^ (shear stress of 4.09e-3 dyn cm^−2^ for the top channel and 0.07 dyn cm^−2^ for the bottom channel). Cell culture media was recirculated, and recirculated media was replaced every alternate day from the falcon tubes. After 4 days of podocyte differentiation, the podocyte induction media was replaced with CultureBoost-R, and the resulting glomerulus chip was maintained for an additional day before treatment with the drugs. The chips were perfused for an additional 2 days with the drug solutions, followed by immunofluorescence analyses of electron microscopy.

For barrier function analyses, CultureBoost-R media was supplemented with 100 μg/mL albumin conjugated to Alexa Fluor 555 (Thermo Fisher Scientific; A34786) and perfused through the bottom channel. Outflow media was collected from the apical channel outlet, and fluorescence intensity was measured using a SpectraMax Fluorescent Plate reader (SpectraMax i3x, Molecular Devices). The amount of Albumin filtered from the basal to the apical channel was analyzed using an equation for renal clearance: ([U] × UV)/[P]), where [U] is the urinary concentration of albumin, UV = volume of media collected from the apical channel outlet reservoir, and [P] = dosing concentration in the basal channel.

For immunofluorescence studies, chips were fixed with 4% paraformaldehyde (Sigma-Aldrich, #16005) in dPBS, followed by permeabilization with 0.125% triton X-100 in dPBS for 5 min. Cells were blocked by incubation with a solution of 1% BSA and 0.125% triton X-100 in dPBS (blocking buffer) for 30 min on ice. The chips were stained with guinea pig Anti-Nephrin (ARP; #GPN02, 1:500), rabbit Anti-Podocin (Abcam, mAb #ab50339, 1:1000), rabbit Anti-TRPC6 (Abcam, pAb #ab228771) and mouse Anti-CD2AP (B-4) (Santa Cruz, #sc-25272) overnight and imaged using the Zeiss 780 Upright confocal microscope equipped with 63x/1.4 Oil Zeiss Plan-Apochromat 44 07 62 (02) WD 0.19 mm objective. Data was analyzed using the Zen 2.3 Black software.

### Transmission electron microscopy

After drug treatment, cells were washed with cold PBS (Gibco; 14190144) followed by fixation with 10 ml of 20% formaldehyde (Sigma-Aldrich, 8.18708), 4 ml of 25% glutaraldehyde (Sigma-Aldrich, G7651), 5 ml of 10× phosphate-buffered saline (Gibco, 14200075), and 31 ml of distilled water (Gibco, 15230001), overnight at 4°C. The chips were postfixed with 1% osmium tetroxide (Electron Microscopy Sciences, 19180) for 1 hour at room temperature in the fume hood, followed by fixation with 0.5% uranyl acetate (Electron Microscopy Sciences, 22400) and dehydration with ethanol (VWR) gradient. The chip’s top compartment was then removed, followed by immersion of the resulting chips in the Spurr’s resin containing 4.1 g of 3,4-Epoxycyclohexanemethyl 3,4-epoxycyclohexanecarboxylate (ERL 4221) (Electron Microscopy Sciences, 15004), 5.9 g of Nonenyl Succinic Anhydride (NSA) (Electron Microscopy Sciences, 19050), 1.43 g of Dow epoxy resins (DER) 736 (Electron Microscopy Sciences, 13000), and 0.1 ml of 2-Dimethylaminoethanol (DMAE) (Electron Microscopy Sciences, 13300), overnight at room temperature. Afterward, the chips were immersed in a freshly prepared Spurr’s resin and cured at 50° to 60°C for 24 to 48 hours. The resulting chip samples were microtome-slice, loaded on TEM grids, counterstained, and visualized using TEM (FEI Tecnai G^2^ Twin).

## SUPPLEMENTARY FIGURES

**Supplementary Figure 1.**
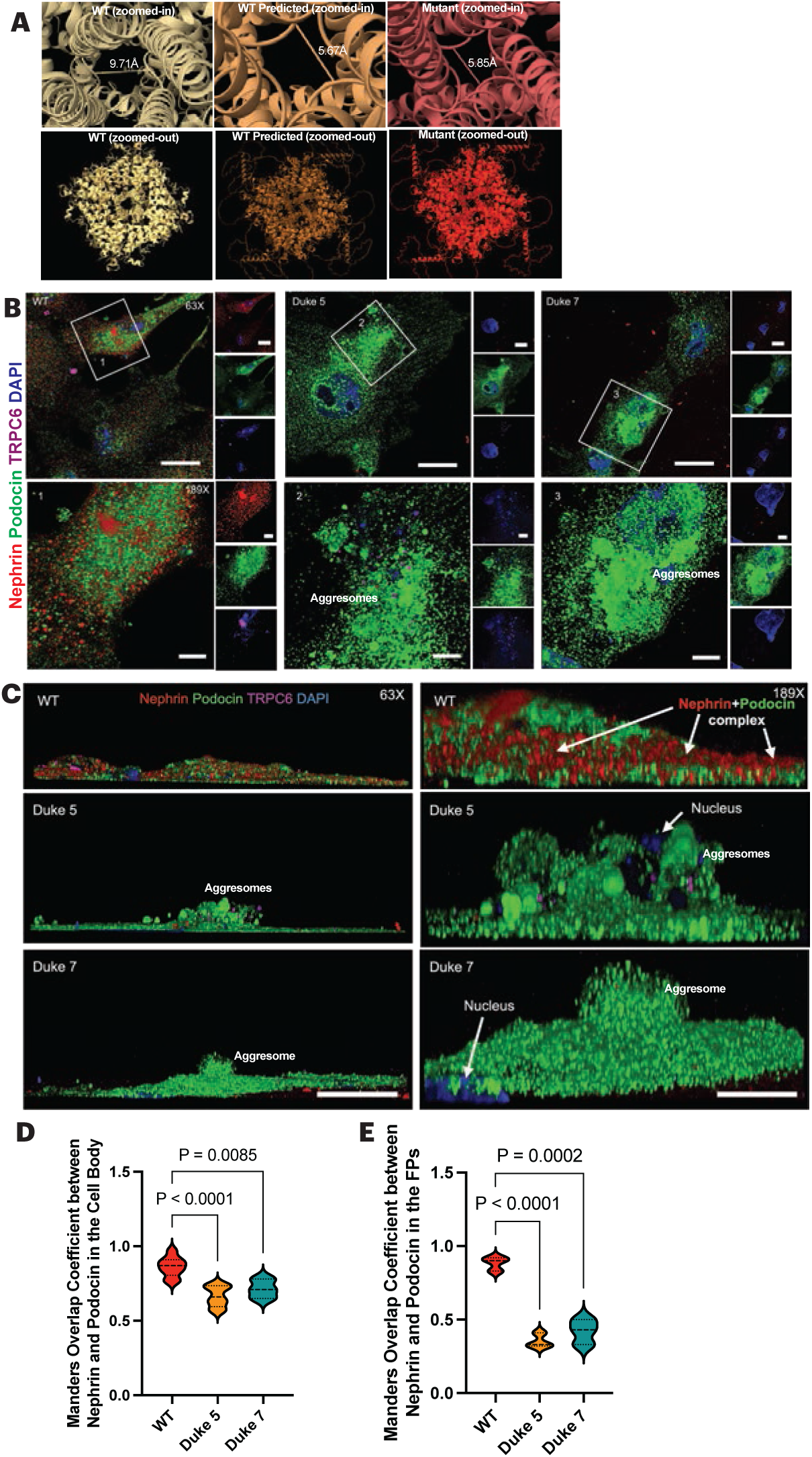
Podocytes differentiated from patient-specific iPS cells demonstrate congestion of slit diaphragm proteins in the endoplasmic reticulum. Panel A shows the pore diameter of the WT, WT predicted, and mutant TRPC6 channel core diameters. Panel B shows the immunofluorescence of protein aggresomes observed in patient-specific iPS cell-derived podocytes’ cell bodies immunolabeled for Nephrin, Podocin, and TRPC6. The bottom panel represents the zoomed-in view. Panel C shows the z-stacks of the podocytes showing Nephrin-Podocin complex formation in the cell body of the podocytes. The mutant podocytes demonstrated massive podocin-enriched aggresomes. Panel D shows Mander’s overlap coefficient for Nephrin and Podocin in the cell body of the untreated podocytes. Panel E shows Mander’s overlap coefficient for Nephrin and Podocin in the foot processes of the untreated podocytes. Panels C and D represent data from three biologically independent experiments, and the data is defined as ± standard error of the mean. We performed one-way ANOVA with Sidak’s multiple comparison post hoc analyses with a family-wise alpha threshold of 0.05. Scale bars: Panel A, top row 20 µm; bottom row 5 µm; Panel B, column 1, 20 µm, column 2, 5 µm.

**Supplementary Figure 2.**
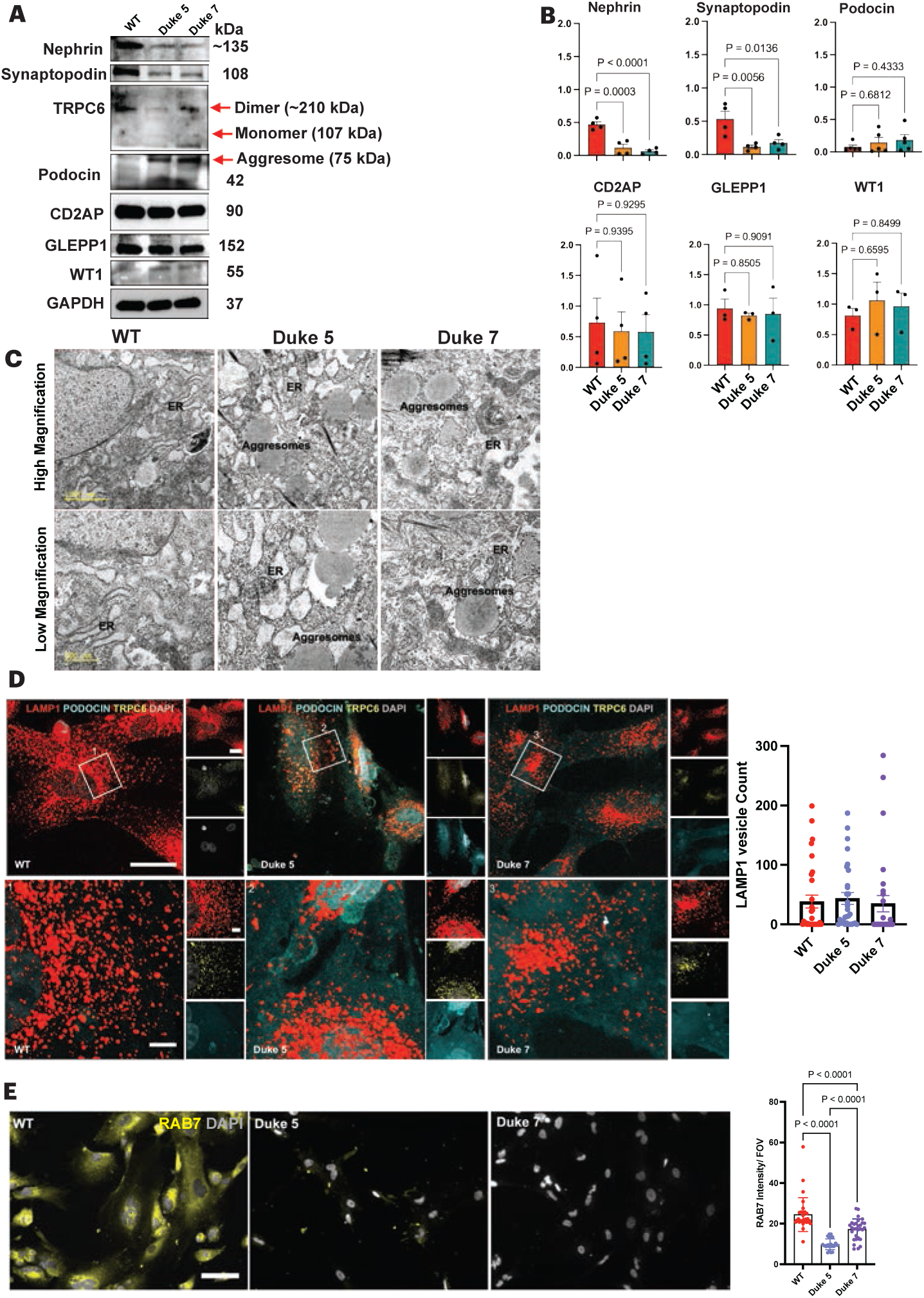
iPS-cell derived podocytes have reduced expression of slit diaphragm proteins, ER congestion, and reduced lysosomes. Panel A shows representative immunoblots of podocyte markers including Nephrin, Synaptopodin, TRPC6, Podocin, CD2AP, GLEPP1, and WT1, the housekeeping protein GAPDH. Panel B shows the quantification of immunoblots for the podocyte markers. Data represents three biologically independent experiments and is defined as ± standard error of the mean. We performed one-way ANOVA with Dunnett’s multiple comparison post hoc analyses with a family-wise alpha threshold of 0.05. Panel C shows electron micrograph of differentiated podocytes with protein aggresomes in the ER of Duke 5 and Duke 7 podocytes. Panel D shows LAMP1 vesicles, Podocin, and TRPC6 expression in podocytes. The LAMP1 vesicle number remains unchanged in the podocytes. Panel E shows lysosomal biogenesis-associated RAB7 expression in the podocytes. RAB7 expression was reduced in the mutants. Data is representative of three biologically independent experiments and the data is defined as ± standard error of the mean. We performed one-way ANOVA with Sidak’s multiple comparison post hoc analyses with a family-wise alpha threshold of 0.05. Scale bar: Panel C, top row 1000 µm, bottom row 500 nm; Panel D, top row 20 µm, bottom row 5 µm; Panel E, 100 µm.

**Supplementary Figure 3.**
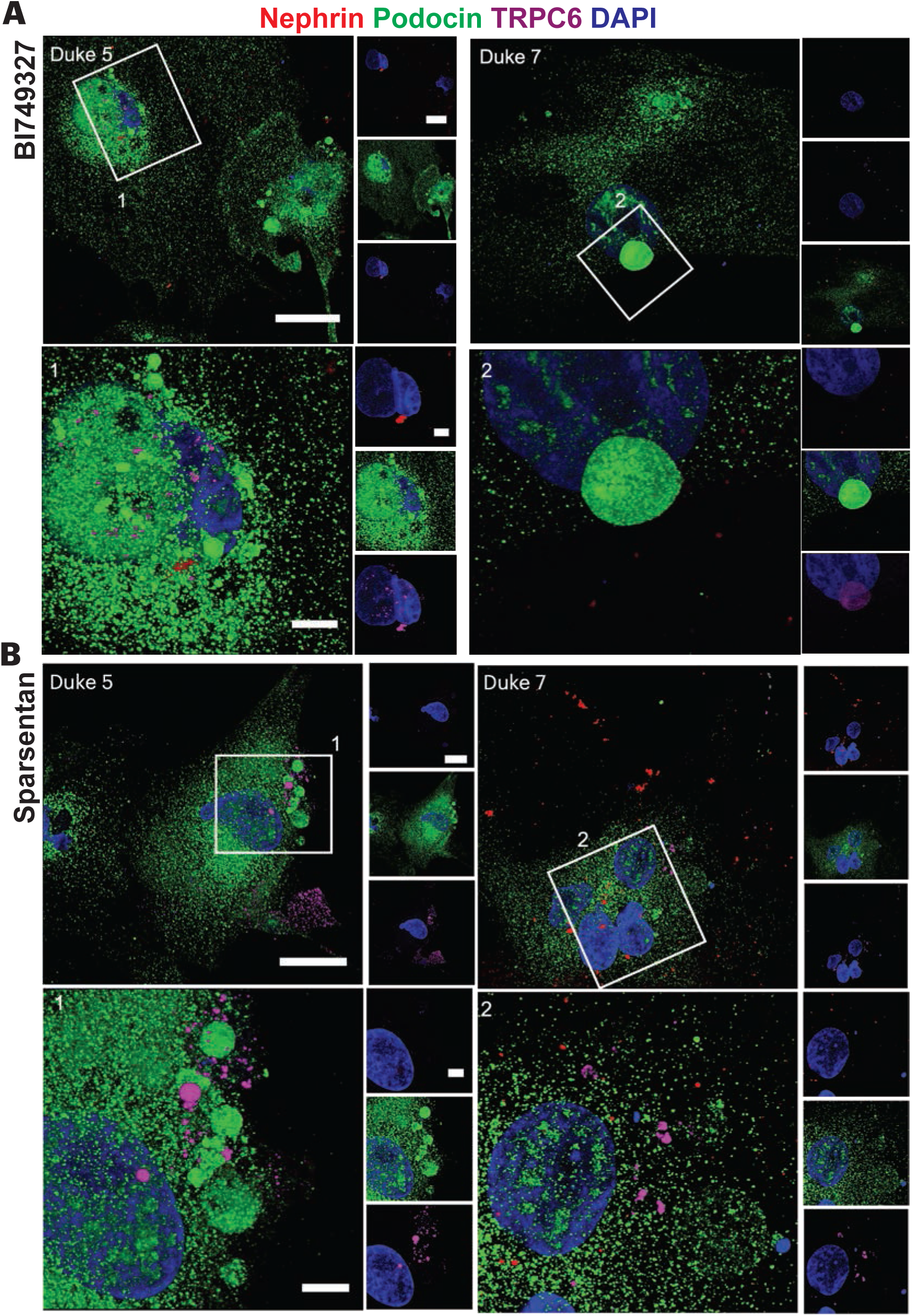
BI749327 and Sparsentan treatment slightly reduced aggresomes but failed to recruit Nephrin. Patient podocytes immunostained for Nephrin, Podocin, and TRPC6 expression after treatment with BI749327 (Panel A) and Sparsentan (Panel B). Scale bars within each panel: top row 20 µm, bottom row 5 µm.

**Supplementary Figure 4.**
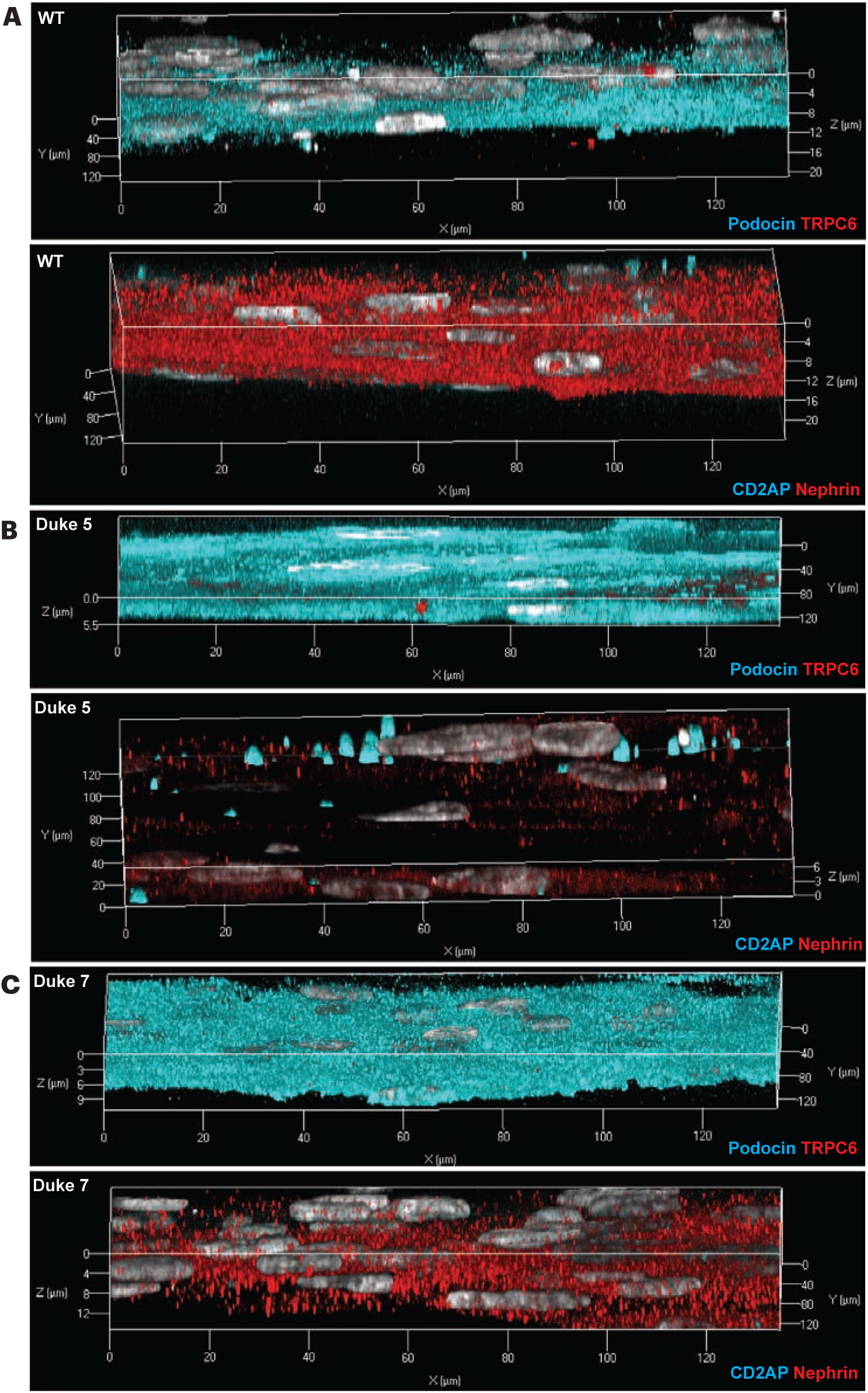
Representative images of patient-specific podocytes differentiated on the glomerular capillary wall on a chip. *Z* stacks of podocytes immunostained for Podocin, TRPC6, Nephrin, and CD2AP in healthy (Panel A) and Duke 5, and C, Duke 7 (Panel B). The *X-, Y-*, and *Z*-axes have been highlighted in the figures.

**Supplementary Figure 5.**
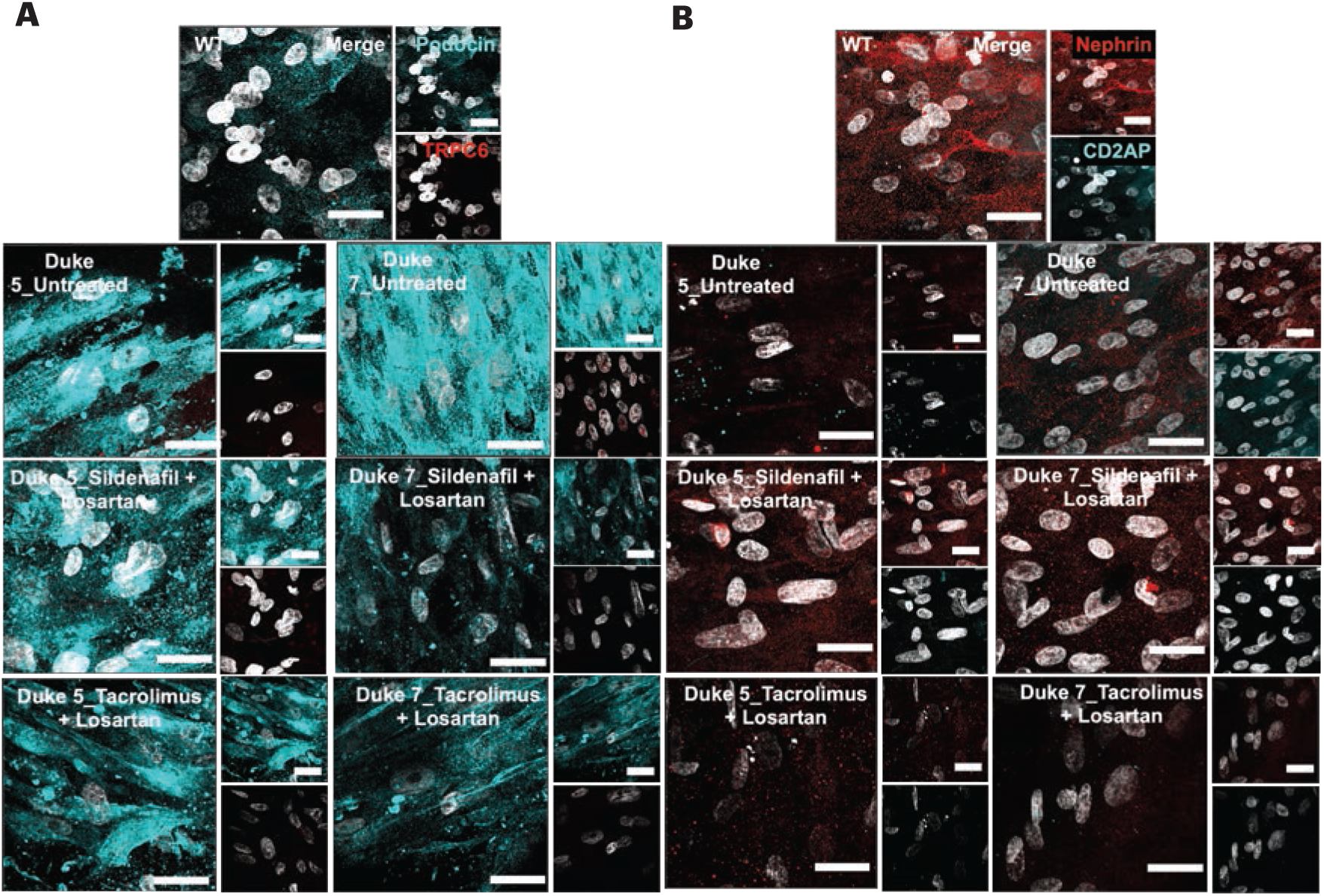
Sildenafil and Losartan treatment reduces podocin aggresomes and recruits Nephrin. Immunofluorescence of podocytes on the glomerular capillary wall-on-a-chip immunostained for Podocin and TRPC6 (Panel A) and Nephrin and CD2AP (Panel B) for WT, Duke 5, and Duke 7 samples. Corresponding quantifications

**Supplementary Figure 6.**
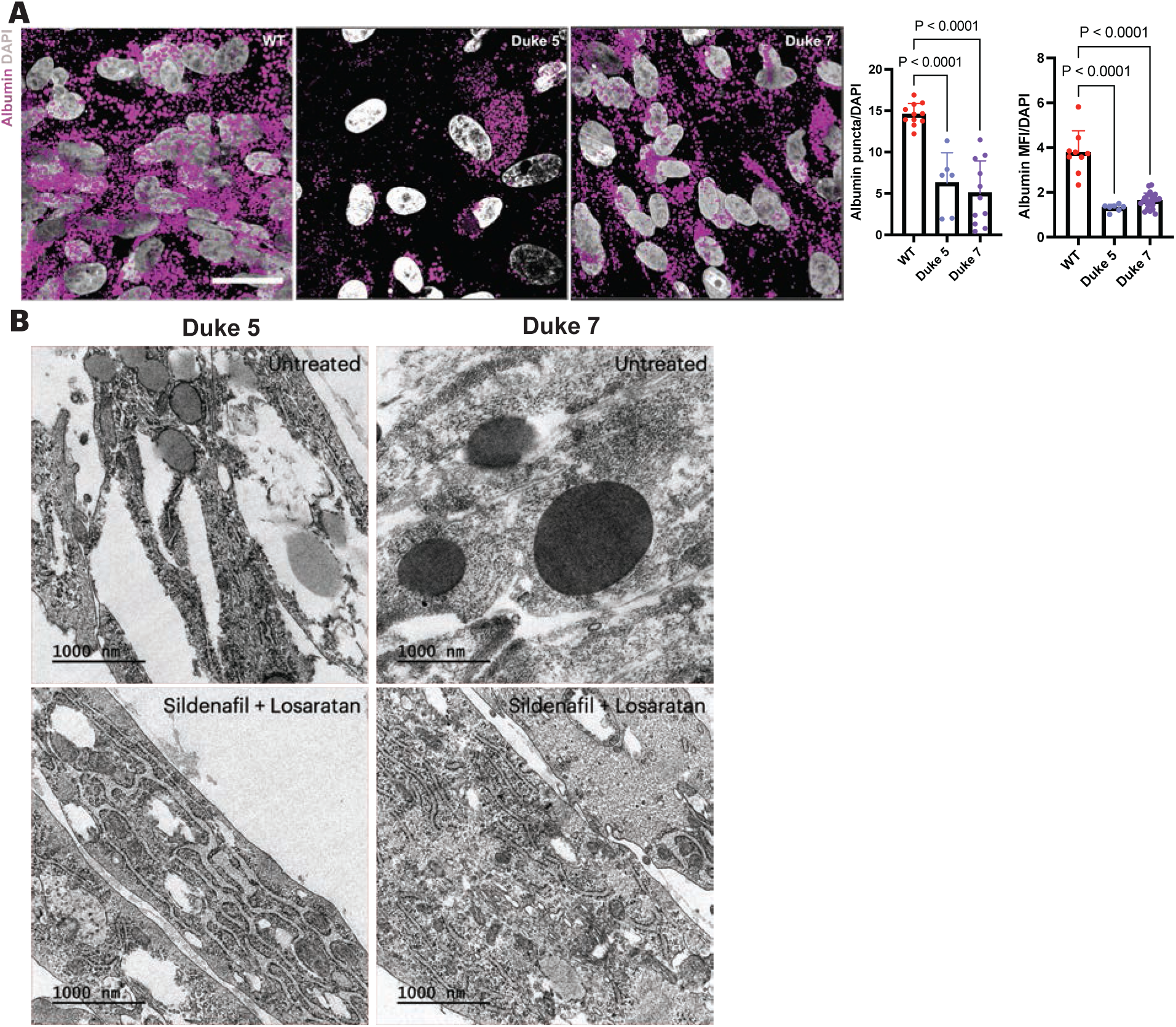
Patient-specific iPS cell-derived podocytes failed to sequester albumin, and Sildenafil and Losartan co-treatment reduced aggresomes in the cell body. Panel A shows albumin sequestration by the podocytes in the glomerular capillary wall-on-a-chip. Albumin uptake was reduced in the mutant podocytes. We performed one-way ANOVA with Dunnett’s multiple comparison post hoc analyses with a family-wise alpha threshold of 0.05. Panel B shows an electron micrograph of untreated and treated podocytes differentiated in the glomerular capillary wall-on-a-chip system. Panel A represents three biologically independent experiments, and the data is defined as ± standard error of the mean. Scale bars: Panel A, 20 µm, Panel B, 1000 nm.

## Notes

### Competing Interest Statement

S. Musah is an inventor on a patent for stem cell differentiation into podocytes and establishment of a glomerulus chip. S. Musah is also an inventor on a pending patent for engineering in vivo-like models of kidney tissues.

## REFERENCE

1. Vivante A. Genetics of Chronic Kidney Disease. N Engl J Med 2024;391(7):627–39.

2. United States Renal Data System. 2022 USRDS Annual Data Report: Epidemiology of kidney disease in the United States. 2022;Available from: https://usrds-adr.niddk.nih.gov/2022

3. Meliambro K, He JC, Campbell KN. Podocyte-targeted therapies — progress and future directions. Nat Rev Nephrol [Internet] 2024 [cited 2024 Jun 30];Available from: https://www.nature.com/articles/s41581-024-00843-z

4. Winn MP, Conlon PJ, Lynn KL, et al. A Mutation in the *TRPC6* Cation Channel Causes Familial Focal Segmental Glomerulosclerosis. Science 2005;308(5729):1801–4.

5. Hall G, Wang L, Spurney RF. TRPC Channels in Proteinuric Kidney Diseases. Cells 2019;9(1):44.

6. Staruschenko A, Ma R, Palygin O, Dryer SE. Ion channels and channelopathies in glomeruli. Physiological Reviews 2023;103(1):787–854.

7. Musah S, Bhattacharya R, Himmelfarb J. Kidney Disease Modeling with Organoids and Organs-on-Chips. Annu Rev Biomed Eng 2024;26(1):annurev-bioeng-072623-044010.

8. Eckel J, Lavin PJ, Finch EA, et al. TRPC6 Enhances Angiotensin II-induced Albuminuria. Journal of the American Society of Nephrology 2011;22(3):526–35.

9. Farmer LK, Rollason R, Whitcomb DJ, et al. TRPC6 Binds to and Activates Calpain, Independent of Its Channel Activity, and Regulates Podocyte Cytoskeleton, Cell Adhesion, and Motility. JASN 2019;30(10):1910–24.

10. Yu H, Kistler A, Faridi MH, et al. Synaptopodin Limits TRPC6 Podocyte Surface Expression and Attenuates Proteinuria. JASN 2016;27(11):3308–19.

11. Riehle M, Büscher AK, Gohlke B-O, et al. TRPC6 G757D Loss-of-Function Mutation Associates with FSGS. JASN 2016;27(9):2771–83.

12. Kuwahara K, Wang Y, McAnally J, et al. TRPC6 fulfills a calcineurin signaling circuit during pathologic cardiac remodeling. J Clin Invest 2006;116(12):3114–26.

13. Farouk SS, Rein JL. The Many Faces of Calcineurin Inhibitor Toxicity—What the FK? Advances in Chronic Kidney Disease 2020;27(1):56–66.

14. Karolin A, Escher G, Rudloff S, Sidler D. Nephrotoxicity of Calcineurin Inhibitors in Kidney Epithelial Cells is Independent of NFAT Signaling. Front Pharmacol 2022;12:789080.

15. Lusco MA, Fogo AB, Najafian B, Alpers CE. AJKD Atlas of Renal Pathology: Calcineurin Inhibitor Nephrotoxicity. American Journal of Kidney Diseases 2017;69(5):e21–2.

16. Musah S, Mammoto A, Ferrante TC, et al. Mature induced-pluripotent-stem-cell-derived human podocytes reconstitute kidney glomerular-capillary-wall function on a chip. Nat Biomed Eng 2017;1(5):0069.

17. Roye Y, Bhattacharya R, Mou X, Zhou Y, Burt MA, Musah S. A Personalized Glomerulus Chip Engineered from Stem Cell-Derived Epithelium and Vascular Endothelium. Micromachines 2021;12(8):967.

18. Mirdita M, Schütze K, Moriwaki Y, Heo L, Ovchinnikov S, Steinegger M. ColabFold: making protein folding accessible to all. Nat Methods 2022;19(6):679–82.

19. Pettersen EF, Goddard TD, Huang CC, et al. UCSF CHIMERAX: Structure visualization for researchers, educators, and developers. Protein Science 2021;30(1):70–82.

20. Mou X, Shah J, Roye Y, Du C, Musah S. An ultrathin membrane mediates tissue-specific morphogenesis and barrier function in a human kidney chip. Sci Adv 2024;10(23):eadn2689.

21. Li SC, Goto NK, Williams KA, Deber CM. Alpha-helical, but not beta-sheet, propensity of proline is determined by peptide environment. Proc Natl Acad Sci USA 1996;93(13):6676– 81.

22. Alías M, Ayuso-Tejedor S, Fernández-Recio J, Cativiela C, Sancho J. Helix propensities of conformationally restricted amino acids. Non-natural substitutes for helix breaking proline and helix forming alanine. Org Biomol Chem 2010;8(4):788.

23. Kocylowski MK, Aypek H, Bildl W, et al. A slit-diaphragm-associated protein network for dynamic control of renal filtration. Nat Commun 2022;13(1):6446.

24. Anderson M, Kim EY, Hagmann H, Benzing T, Dryer SE. Opposing effects of podocin on the gating of podocyte TRPC6 channels evoked by membrane stretch or diacylglycerol. American Journal of Physiology-Cell Physiology 2013;305(3):C276–89.

25. Wang Q, Tian X, Wang Y, et al. Role of Transient Receptor Potential Canonical Channel 6 (TRPC6) in Diabetic Kidney Disease by Regulating Podocyte Actin Cytoskeleton Rearrangement. Journal of Diabetes Research 2020;2020:1–11.

26. Lin BL, Matera D, Doerner JF, et al. In vivo selective inhibition of TRPC6 by antagonist BI 749327 ameliorates fibrosis and dysfunction in cardiac and renal disease. Proc Natl Acad Sci USA 2019;116(20):10156–61.

27. Kohan DE, Bedard PW, Jenkinson C, Hendry B, Komers R. Mechanism of protective actions of sparsentan in the kidney: lessons from studies in models of chronic kidney disease. Clinical Science 2024;138(11):645–62.

28. Rheault MN, Alpers CE, Barratt J, et al. Sparsentan versus Irbesartan in Focal Segmental Glomerulosclerosis. N Engl J Med 2023;NEJMoa2308550.

29. Hall G, Rowell J, Farinelli F, et al. Phosphodiesterase 5 inhibition ameliorates angiontensin II-induced podocyte dysmotility via the protein kinase G-mediated downregulation of TRPC6 activity. American Journal of Physiology-Renal Physiology 2014;306(12):F1442– 50.

30. Sonneveld R, Hoenderop JG, Isidori AM, et al. Sildenafil Prevents Podocyte Injury via PPAR-γ–Mediated TRPC6 Inhibition. JASN 2017;28(5):1491–505.

31. Pușcașu C, Zanfirescu A, Negreș S, Șeremet OC. Exploring the Multifaceted Potential of Sildenafil in Medicine. Medicina 2023;59(12):2190.

32. Benzing T, Salant D. Insights into Glomerular Filtration and Albuminuria. N Engl J Med 2021;384(15):1437–46.

33. Ahmed WS, Geethakumari AM, Biswas KH. Phosphodiesterase 5 (PDE5): Structure-function regulation and therapeutic applications of inhibitors. Biomedicine & Pharmacotherapy 2021;134:111128.

34. Pearce LR, Komander D, Alessi DR. The nuts and bolts of AGC protein kinases. Nat Rev Mol Cell Biol 2010;11(1):9–22.

35. Ranek MJ, Terpstra EJM, Li J, Kass DA, Wang X. Protein Kinase G Positively Regulates Proteasome-Mediated Degradation of Misfolded Proteins. Circulation 2013;128(4):365–76.

36. Ranek MJ, Kokkonen-Simon KM, Chen A, et al. PKG1-modified TSC2 regulates mTORC1 activity to counter adverse cardiac stress. Nature 2019;566(7743):264–9.

37. Moller CC, Wei C, Altintas MM, et al. Induction of TRPC6 Channel in Acquired Forms of Proteinuric Kidney Disease. Journal of the American Society of Nephrology 2007;18(1):29– 36.

